# MCX Cloud – a modern, scalable, high-performance and in-browser Monte Carlo simulation platform with cloud computing

**DOI:** 10.1101/2021.06.28.450034

**Authors:** Qianqian Fang, Shijie Yan

## Abstract

**Significance:** Despite the ample progress made towards faster and more accurate Monte Carlo (MC) simulation tools over the past decade, the limited usability and accessibility of these advanced modeling tools remain key barriers towards widespread use among the broad user community.

**Aim:** An open-source, high-performance, web-based MC simulator that builds upon modern cloud computing architectures is highly desirable to deliver state-of-the-art MC simulations and hardware acceleration to general users without the need for special hardware installation and optimization.

**Approach:** We have developed a configuration-free, in-browser 3-D MC simulation platform – MCX Cloud – built upon an array of robust and modern technologies, including a Docker Swarm-based cloud-computing backend and a web-based graphical user interface (GUI) that supports in-browser 3-D visualization, asynchronous data communication, and automatic data validation via JavaScript Object Notation (JSON) schemas.

**Results:** The front-end of the MCX Cloud platform offers an intuitive simulation design, fast 3-D data rendering, and convenient simulation sharing. The Docker Swarm container orchestration backend is highly scalable and can support high-demand GPU MC simulations using Monte Carlo eXtreme (MCX) over a dynamically expandable virtual cluster.

**Conclusion:** MCX Cloud makes fast, scalable, and feature-rich MC simulations readily available to all biophotonics researchers without overhead. It is fully open-source and can be freely accessed at http://mcx.space/cloud.

## 1 Introduction

Since the initial release of the first open-source Monte Carlo (MC) light transport simulator – MCML^1^ – nearly 30 years ago, MC-based photon simulations have been playing important roles amongst the biophotonics research community to facilitate the design and optimization of novel imaging instrumentation and image reconstruction, as well as providing gold-standard solutions for validating novel algorithms and data analysis pipelines. Notably, in the last decade, a list of free and open-source MC simulators have been published and further improved upon by their respective authors. The proliferation of open-source MC tools provides the community with abundant options to meet diverse needs arising in biophotonics research.

Many of the emerging MC simulators have placed strong emphases towards addressing two of the top limitations facing traditional MC algorithms. First, the adoption of massively parallel computing and graphics processing units (GPUs) have greatly improved the computational efficiency of conventional MC simulations, shortening the simulation run-time by tens to hundreds fold on a modern GPU.^2–7^ In parallel, a list of new MC algorithms were proposed to handle more complex and accurate tissue anatomical boundaries.^8–11^ Among these algorithms, mesh-based Monte Carlo (MMC) offers the capability to accurately model a curved tissue boundary with tetrahedral meshes while performing ray-tracing computation significantly more efficiently than surface-based MC techniques.^8^ More recently, hybrid approaches that combine shape representations offer further computational efficiency and accuracy.^12–15^ These hybrid approaches include 1) dual-grid MMC (DMMC)^12^ that combines a coarse tetrahedral mesh with a dense voxelated output volume, 2) split-voxel MC (SVMC)^14^ that combines curved surface meshes within a compact voxel data structure, and 3) implicit MMC (iMMC)^15^ that combines a skeletal tetrahedral mesh with implicitly defined shapes such as tubes, spheres and thin membranes. These enhancements in modeling geometry have resulted in significantly improved accuracy, which can be directly translated to further speed enhancement while achieving the same output accuracy as conventional approaches.

Compared to many published traditional research codes that were developed as single-release static software, an increasing number of new MC software packages have started tackling the challenges of usability and long-term maintainability. Many of these projects openly embrace state-of-the-art software engineering best practices and offer the software as a vibrantly growing platform via continuous enhancements, timely bug fixes, and active user support via flexible feedback channels. Ease-of-use has also become the focus of a number of recently published MC toolkits, where MATLAB-based dynamic library (MEX) interfaces and graphical user interfaces (GUIs) have been reported.^16, 17^

With the exciting progress in developing open-source MC simulators with increasing speed, functionality, accuracy, and user-friendliness, we would like to tackle here the next major challenge in high-performance, general-purpose MC photon simulation software, namely scalability and availability. A number of previous publications, including several from our group, have addressed the challenges in creating scalable simulations that can utilize more than one GPU or run simulations across CPUs/GPUs of multiple vendors. In particular, a number of previous papers reported OpenCL-based MC implementations^5, 18^ that are readily scalable across heterogeneous computing environments including multi-vendor hardware. Several NVIDIA CUDA-based GPU MC simulators also offer support to multiple GPU architecture generations and multi-GPU simulations. Regarding availability, most MC software tools are disseminated via the conventional download-installation-execution approach. Software dissemination via Docker-based container images has also become increasingly popular and is found in several notable open-source MC tools, including MCX,^3^ MMC^10^ and FullMonte.^19^ Nevertheless, a majority of these software dissemination methods require users to have a pre-configured GPU to be able to execute their desired simulations. Purchasing and configuring high-performance GPUs may still present a barrier for beginner and less-experienced computer users. Online-based MC modeling tools that do not require local GPU installation are extremely limited. In 2011, a proprietary web-based MC simulator, MCOnline,^20^ was reported by Doronin and Meglinski using Microsoft Silverlight and ASP.NET technologies as the front-end and a GPU MC simulator on the server-side. Although this tool is still being actively maintained, the proprietary nature of the tool and the limited scalability of the underlying technologies necessitate a re-investigation using up-to-date cloud-computing technologies. In 2020, another proprietary web-based MC simulation platform, Multi-Scattering, was published by Joönsson and Berrocal,^21^ featuring a modern and user-friendly web GUI design, versatile scattering phase function support, and a proprietary voxel-based MC simulator in the backend. While this tool offers intuitive interfaces to attract a broad userbase, the maximum simulation domain is limited to 20 *×* 20 *×* 20 voxels,^21^ making it quite limited for solving practical problems.

In this work, we report a modern, scalable, high-performance, and fully open-source in-browser MC simulation platform – MCX Cloud – to bring state-of-the-art GPU hardware and our extensively-optimized and feature-rich MCX simulator software to the rapidly growing biophotonics research community. Our MCX Cloud platform embraces an array of modern and standardized cloud-computing techniques. In the backend, it utilizes Docker (https://docker.com/) and Docker Swarm-based container orchestration technology to create a highly scalable, dynamically expandable, fault-tolerant, and distributed GPU virtual cluster with built-in “ingress load-balancing” capabilities. In the front-end, we have developed a modern web GUI based upon a list of open-source web technologies, such as HTML5 markup language,^22^ cascading style sheets (CSS), JavaScript, and JQuery (https://jquery.com/) for GUI development, and WebGL^23^ for in-browser 3-D data rendering.

A key advancement that enables us to develop such a compact, scalable and portable software/hardware platform is the adoption of JavaScript Object Notation (JSON, https://json.org) and JData – an open-specification for scientific data annotation using JSON^24^ – as the input and output data formats for MCX. JSON is a lightweight, human-readable, and ubiquitously supported data format that is capable of storing complex hierarchical data. It has rapidly replaced XML (extensible markup language) as become one of the most widely used data exchange formats among web applications. Since 2012, we have migrated MCX’s input file format to JSON and subsequently completed the migration of all output data files to JSON in 2020. In this work, we use JSON Schema^25^ – an open-standard for defining JSON-based data files – and JSON Editor – a lightweight JavaScript library for editing arbitrary JSON files inside a browser – to create a compact and easy-to-maintain in-browser MCX input editor and data visualization platform that is intuitive to use for users without any programming experience. Both the front-end and backend designs in MCX Cloud are highly flexible and require only minimal changes to support additional input/output fields and hardware extensions. In contrast with previously published online MC simulators, both the front-end (user interface) and the backend (server-side scripts) of MCX Cloud are open-source so that a user may easily configure a private cloud-computing virtual cluster to run MCX-based simulations from a browser.

In the following sections, we will first discuss the key technology components that have enabled this scalable cloud-computing based MC simulator, including a brief discussion on the latest MCX light transport simulator, backend design, front-end design, and input/output data formats. We then show a number of example simulations and a benchmark demonstrating scalability for high-performance, distributed GPU-based simulations using MCX Cloud. Finally, we discuss our plans for further improvement of this platform.

## 2 Methods

A diagram showing the overall design of the MCX Cloud simulation platform is shown in Fig. 1. This highly portable and scalable platform can be divided into a front-end (web-based user interface) and a backend (a distributed GPU cluster managed by Docker Swarm services), communicating asynchronously via lightweight and versatile JSON/JData data packets. The key technologies used in this platform are highlighted in gray-shaded boxes, and open-source software/libraries used are highlighted in orange colored text. The MCX Docker image (bottom-left) – a lightweight package that contains the MCX simulator software along with all dependencies – is publicly hosted on Dockerhub. In the following subsections, we will discuss each key component and the overall simulation workflow.

**Fig 1.**
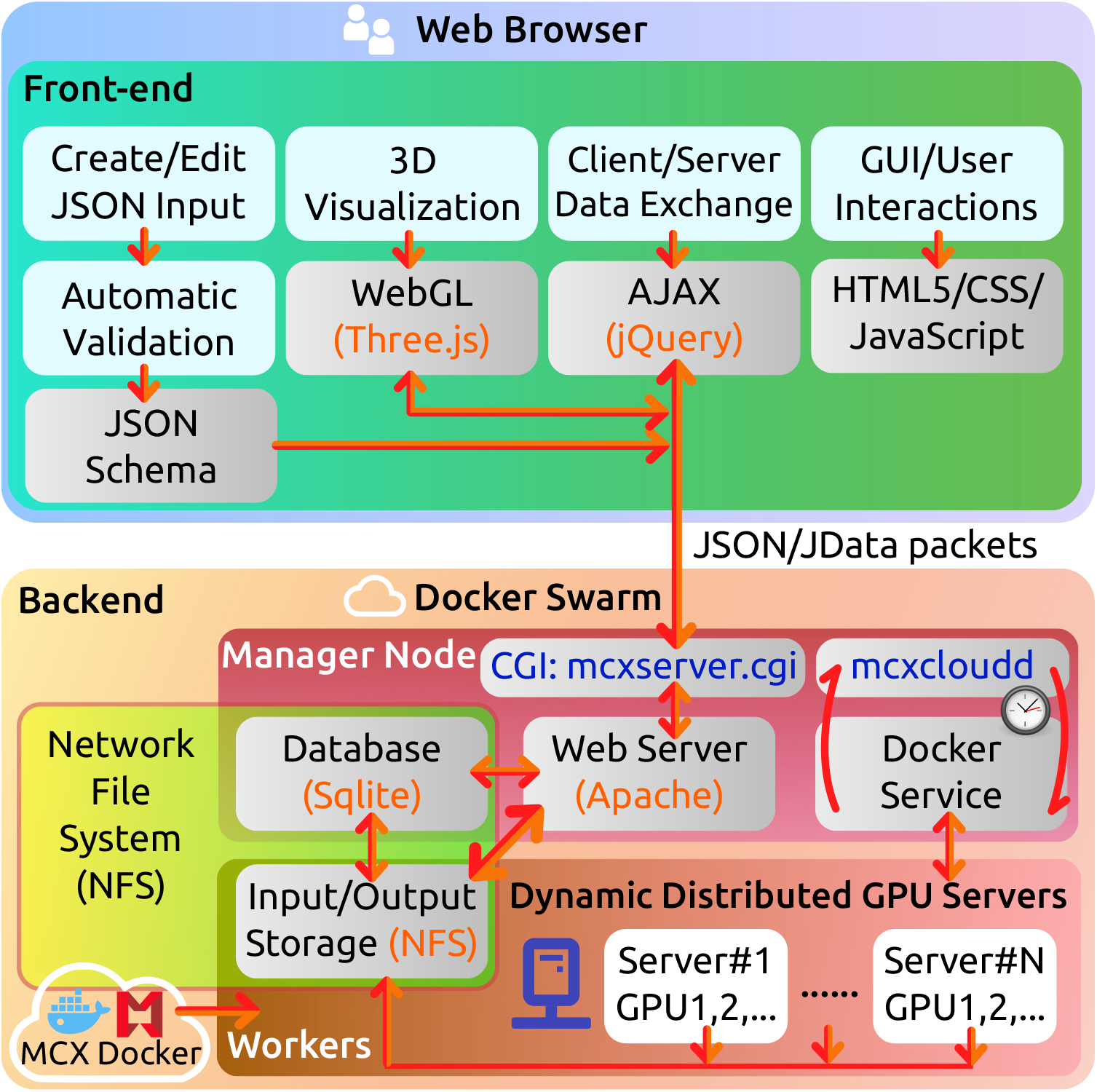
Diagram showing the overall design of the MCX Cloud simulation platform. Gray-shaded boxes indicate key technologies utilized in this platform; boxes shaded in light-blue indicate key functionalities.

### 2.1 MCX photon transport simulator and containerization

At the heart of this cloud computing platform is a Docker container image of our latest MCX photon simulator. A container is simply a lightweight package that allows users to reliably reproduce the virtual environment, including dependencies and libraries, of a given application and conveniently execute it consistently across various platforms. A containerized application automatically downloads all dependencies necessary to run the program, greatly simplifying the installation and configuration process of new software. In this work, our MCX container image is built using the “base image” `cuda-9.0` provided by NVIDIA and is publicly accessible via Dockerhub – one of the largest repositories of container images.

The current release of the MCX photon simulator contains numerous algorithmic improvements over the original version published in 2009.^3^ Briefly, MCX is a GPU-accelerated, parallel MC photon transport simulator that supports 3-D heterogeneous media defined in a voxelated space. We want to particularly highlight several key improvements over the original MCX algorithm described in Fang *et al*.^3^ First, we have implemented precise ray-tracing in MCX releases since 2016. Photon trajectories are precisely broken into segments bounded by voxel boundaries; in comparison, the original MCX accumulates photon energy at a fixed 1-mm spacing along the trajectory. This update has led to significant accuracy improvements in simulation results. Secondly, all MCX releases since 2013 have supported over a dozen complex source types, including pencil beam, isotropic source, planar and disk sources, Gaussian beam, Fourier patterns (for spatial-frequency domain imaging, or SFDI), line and slit sources, user-defined 2-D and 3-D pattern sources, etc. For all area-sources, a focal-length parameter is also added to enable convergent and divergent beams. Thirdly, four new boundary conditions (BCs) are supported on the bounding box of the voxelated domain, including a total absorption BC, a Fresnel reflection BC, a total reflection/mirror BC, and a cyclic BC (photons exiting from a bounding box face re-enters from the opposite face to simulate infinite medium). Fourthly, MCX outputs a variety of detected photon data outputs, including partial-pathlengths, partial-scattering-event-count, exiting position and direction, momentum transfer, initial photon weight etc. Moreover, MCX not only supports label-based segmented volume, but also continuously varying medium. Furthermore, MCX has incorporated state-of-the-art MC algorithm advances, including photon replay,^26^ photon sharing,^12^ and our latest hybrid algorithm split-voxel MC (SVMC).^14^ Lastly, we have extensively optimized the MCX GPU computing implementation and dramatically improved its simulation speed across multiple generations of NVIDIA GPU architectures. We want to highlight that MCX is an actively maintained platform funded by the National Institute of Health (NIH). New features are constantly being added; recently added key features include user-defined scattering phase functions and modeling of polarized light in 3D heterogeneous media.

### 2.2 Docker Swarm based cloud computing backend

Docker Swarm is a lightweight container “orchestration” framework that is built-in to the Docker toolkit. Docker Swarm allows users to create a virtual cluster made of a single or multiple Docker service “nodes”, dispatch executions across such distributed computing environments, and perform job distribution and job queue management. In our current MCX Cloud configuration, we have included several rack-mount servers as Docker service nodes and also enumerated each GPU hosted on each server as a named resource. As a result, any simulation dispatched by the Docker service to the Swarm can be automatically assigned to one of the vacant GPU cards among all participating nodes, determined automatically by the Docker Swarm manager node. Utilizing the Docker Swarm framework to manage the computing hardware backend offers a number of notable benefits. First, a Docker Swarm can be dynamically expanded and shrunk without interrupting current jobs. Therefore, system administrators can grow the number of GPUs to accommodate the job loads or shutdown some of the nodes for maintenance without interrupting the simulation queue. Secondly, the latest Docker Swarm release offers fine-grained GPU-based resource allocation and job distribution capability. With a simple configuration, one can let Docker Swarm assign each simulation to a single GPU or to a single host, utilizing all GPUs on the host in parallel. The Docker Swarm platform also provides high fault tolerance: when a hardware failure is detected on a host or a GPU, incomplete jobs can be automatically relaunched by the Docker service manager. We would like to emphasize that the Docker platform is a vastly rich ecosystem for cloud computing; numerous free tools are available for container creation, sharing, management, and orchestration. In this initial release of MCX Cloud, we chose Docker Swarm as the orchestration framework largely because of its simplicity, but our platform can be further adapted to support other orchestration platforms such as Kubernetes or Apache Mesos.

### 2.3 JSON/JData based data exchange format and JSON Schema

As we mentioned previously, JSON is an internationally standardized (also known as ISO21778:2017) data exchange format, and is at the core of most today’s web-based applications. Compared to XML, JSON is extremely lightweight and fast to parse, yet it is capable of storing complex hierarchical data. Numerous free and lightweight JSON parsers are available today for nearly all existing programming languages, permitting plug-and-play implementation of JSON data support in most applications.

Despite these aforementioned advantages, adoption of JSON in storing scientific data is largely limited to handling lightweight metadata. This is because JSON does not have explicit rules on how to serialize common scientific data structures such as N-dimensional (N-D) arrays, complex and sparse arrays, tables, graphs, trees, etc. Additionally, JSON does not directly support storage of strong-typed binary data. To bridge this gap, our group published an open-standard – the JData Specification^24^ – to systematically serialize common data structures used in scientific research, enabling storage of binary strongly-typed data using 100% JSON-compatible annotation tags. In addition, the JData specification also provides a binary data interface utilizing the Universal Binary JSON (UBJSON, https://ubjson.org) format to offer additional space efficiency and processing speed. In Listing 1, we show an input data file snippet that MCX uses to define an MCX simulation. In the `“Shapes”` section, an example defining a 3-D volume using the JData annotations is shown.

In addition to using JSON to encode input data, we have also completed the migration of MCX volumetric output data as of 2020, converting from the NIfTI data format^27^ to JSON/JData-based JNIfTI^28^ data files. Additional output data associated with detected photon data, including partial pathlengths and exiting position, are also stored in a JSON/JData^24^ file that is readily readable by any existing JSON parser. The migration from a opaque and rigid binary conventional format to the human-readable and easily extensible JSON/JData file sets the foundation for migrating MCX from a local application to the cloud and web environments.

A key benefit of adopting JSON based data formats is to enable machine-automatable data validation. This can be readily achieved using the JSON Schema framework. JSON Schema is a systematic approach to defining data types, formats, and properties for each data entry in a JSON data structure, and is currently a proposed Internet standard by the Internet Engineering Task Force (IETF).^25^ It has received widespread adoption for automating and creating JSON based data files. In this work, we have rigorously defined the JSON-based MCX input file format using JSON Schema syntax (which is fully JSON-compatible). A snippet of MCX input file JSON schema is shown in Listing 2.

**Listing 1.**
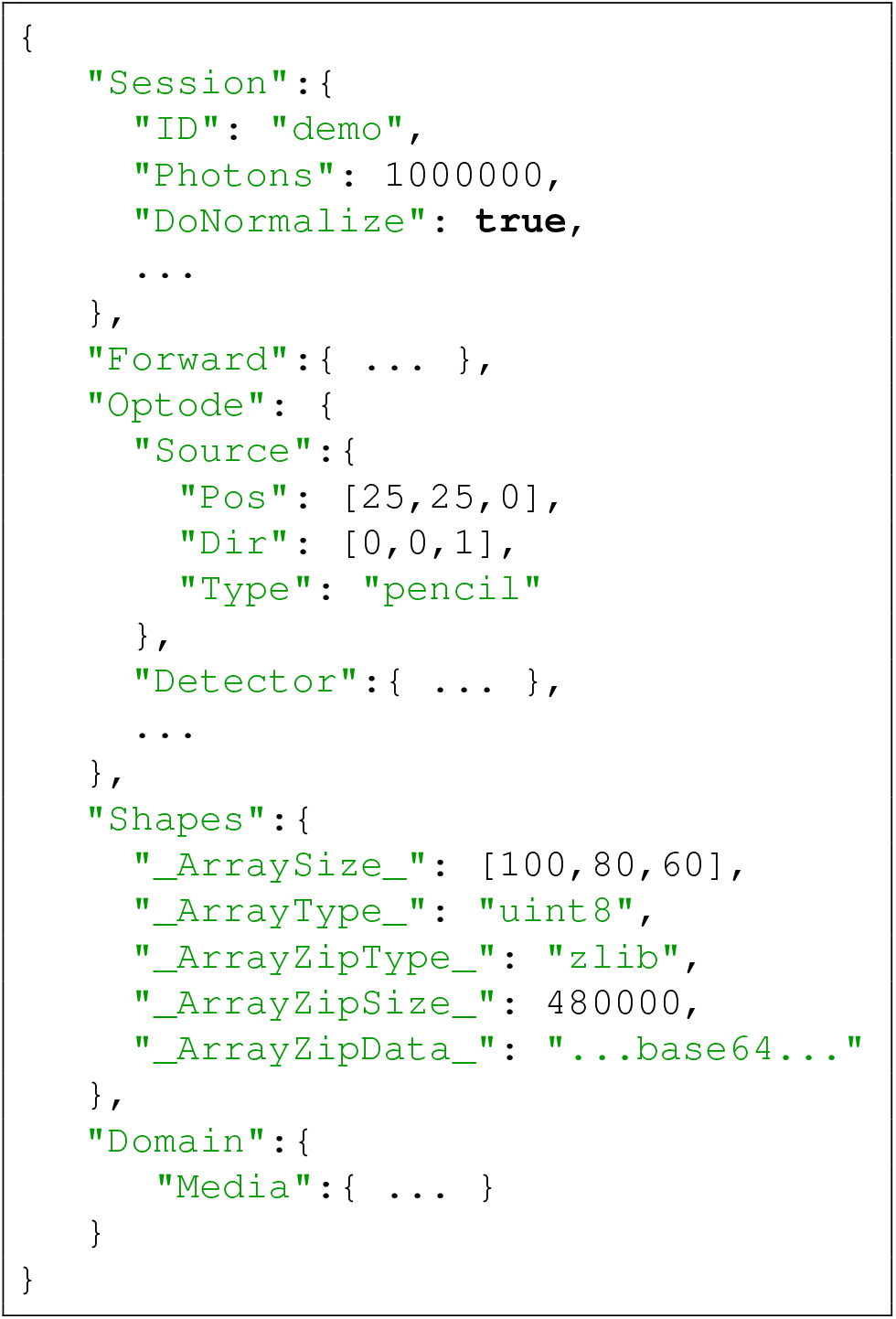
JSON based MCX input file

**Listing 2.**
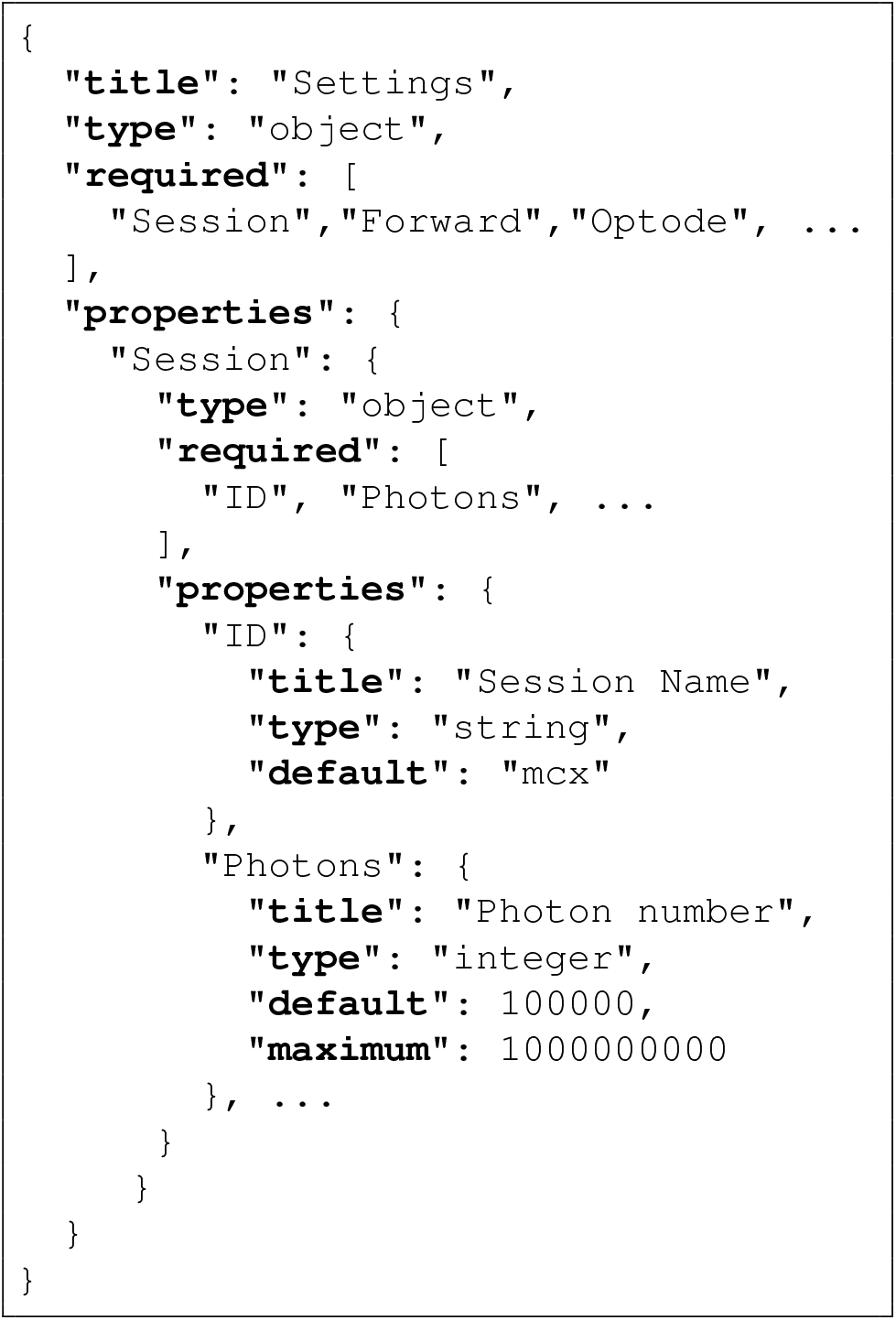
JSON Schema sample

### 2.4 Web-based JSON editor and graphical user interface design

The front-end, i.e. the web GUI, of MCX Cloud consists of two major components – an in-browser JSON data editor to create JSON-formatted input data for MCX simulations and a 3-D data rendering module based on WebGL (see below section). The web-based MCX JSON input editor was derived by combining an open-source general-purpose JSON editor developed by Jeremy Dorn *et al*. and our JSON-schema of MCX input JSON data format. The JSON editor module is a lightweight (73 kB in size) JavaScript library that enables the creation and editing of arbitrary JSON-formatted data using a user-defined schema. It also simultaneously supports a number of popular web GUI frameworks icon libraries to improve customizability.

A minimalistic design style is used to provide users with a clean and streamlined environment to create, preview, execute, render, and easily share MCX simulations. All front-end functionalities are achieved using a combination of HTML5 and JavaScript programming. Notably, the use of the JQuery library makes the front-end compact (less than 1,500 lines of JavaScript code) and easy-to-maintain.

### 2.5 In-browser rendering of 3-D shapes and volumetric data using WebGL

In the front-end of MCX Cloud, we have developed fully-featured 3-D shape and volumetric data rendering and download functionalities. In comparison, the web GUI of MCOnline only provides rendering and data downloading for a particular x/y/z slice of the volume. The in-browser 3-D data rendering feature is enabled by the WebGL technology,^23^ conveniently provided via utilizing the open-source Three.js JavaScript rendering library (https://threejs.org) application programming interfaces (APIs).

Our MCX JSON input file accepts two methods for defining a heterogeneous simulation domain: 1) a constructive solid geometry (CSG) approach using a list of shape primitive constructs such as spheres, boxes, cylinders, x/y/z layered structures etc, and 2) a JData-formatted^24^ 3-D array that defines the tissue-types or per-voxel absorption/scattering values of a voxelated space. As a result, in our web GUI, we support rendering of both shape-based domain configurations as well as 3-D array based rendering. An OpenGL 3-D texture is created if a 3-D array-based volume is provided; the voxelated input domain is rendered in either maximum-intensity-projection (MIP) or isosurfaces. In either case, convenient controls of 3-D rotation and zooming are supported. Because Three.js is highly optimized on modern browsers such as Chrome and Firefox, rendering a typically sized volume does not noticeably increase the CPU/GPU loads of the browser.

### 2.6 Asynchronous data communication and optimization

The client (i.e. web GUI) and the server (i.e. a web service running on the manager node of the Docker Swarm) communicate via asynchronous data communication, known as AJAX (asynchronous JavaScript And XML). Despite the name, JSON, instead of XML, has been predominantly used in today’s web applications data exchange. User inputs are encoded as lightweight JSONP (JSON with Padding) data packets and sent to the server; the server sends back the response, also encoded as JSON packets, and informs the JavaScript on the web GUI to update the web page content dynamically without needing to reload the entire web page.

To facilitate the processing of user submissions and management of Docker Swarm jobs, we developed an ultra-compact CGI (common gateway interface) script, named “mcxserver”, written in the Perl programming language to handle user submitted job requests. These submitted simulation data are stored in a database using Sqlite (https://sqlite.org) for fast query and update. The “mcxserver” server script also handles status queries from the client once a job is submitted, and returns the simulation output data once the simulation is completed. A random hex-key is assigned to each submitted job to uniquely identify a given job. In addition, another Perl script named “mcxcloudd” (MCX Cloud Daemon, see Fig. 1) is repeatedly executed at a fixed time interval (currently set to run every 20 seconds) and checks 1) if the Docker Swamp has a vacant GPU device, and 2) if there exists unprocessed user-submitted job requests in the Sqlite database. If both are confirmed, a `docker service` command is then submitted to launch the user submitted job to the Docker Swarm. The web server database and simulation input/output files are shared among all Docker Swarm nodes via the network file-system (NFS), as depicted in Fig. 1.

To optimize server disk usage, we define a job expiration time window (currently set to 1 hour) and configure another recurrent process (known as a “cron-job”) to automatically clean the expired job folders to save space. If a simulation is frequently executed by users, such as the default simulation or built-in examples, we keep the simulation output folder in a cache folder to avoid repeated computation.

### 2.7 Reusable and community-driven simulation repository

Guided by the FAIR principle^29^ (i.e. making data findable, accessible, interoperable and reusable), our MCX Cloud platform provides convenient mechanisms to allow a user to share their simulations with the community or reuse simulations contributed by other users. In MCX Cloud’s “Share” tab, a user can fill out a simple form to give permission for others to use his/her designed MCX simulation JSON data. A dedicated server database is used to store these shared simulation settings. When a user opens the “Browse” tab in the web GUI, the GUI retrieves a list of community-contributed simulations, including the JSON input data as well as a domain preview thumbnail. If a user clicks on any one of the previously defined simulations, the JSON data corresponding to the selected simulation will be loaded and ready for modification by the user. Over time, we anticipate that this feature will eventually build a rich simulation repository, not only helping new users quickly create new and more advanced simulations, but also establish a set of standardized benchmarks that facilitate cross-validation between diverse light simulation tools.

## 3 Results

Following the methodologies discussed above, we have created a preview version of the MCX Cloud simulation platform. In this initial configuration of the MCX Cloud backend, we have currently included 6*×* Docker service nodes using 6 Linux servers running Ubuntu 16.04 and 20.04 and Docker version 20.10.3. To balance the server loads, one of the servers is configured as the “manager” node and is dedicated to running the web service (Apache 2.4.18), the CGI script (“mcxserver”) and the “mcxcloudd” cron-job to process the user-submitted job queue, as shown in Fig. 1. The remaining servers host a total of 5*×* NVIDIA RTX 2080 SUPER (Turing) GPUs, 4*×* GTX 1080 (Pascal) GPUs, and 1*×* GTX 980Ti (Maxwell) GPU. This preview Docker Swarm backend is capable of simultaneously executing 10 parallel simulations. With only a few simple commands, we can effortlessly expand this Docker Swarm to include more nodes and GPUs without interrupting the service. Docker provides command-line tools to enable easy administration of the Docker Swarm and the jobs running on it. Graphical management tools are also freely available, including Portainer and Shipyard.

To demonstrate the GUI design in MCX Cloud’s front-end, in Fig. 2, we include four screenshots showing (a) the main menu screen, (b) the “Browse” tab for loading built-in or community-contributed simulation library, (c) the “Create” tab for MCX input JSON data in-browser editing, and (d) the “Run” tab for job submission and management. The initial loading of the front-end web GUI only needs to download a total of 570 kB of resources, including 9 open-source JavaScript libraries, two cascade style-sheets (CSS), three web-fonts, and a single HTML file. Small software footprint enables smooth access to this cloud service even for users with low-bandwidth networks. All subsequent data exchange with the server is achieved via AJAX with lightweight JSON data packets; no web page reloading is needed.

**Fig 2.**
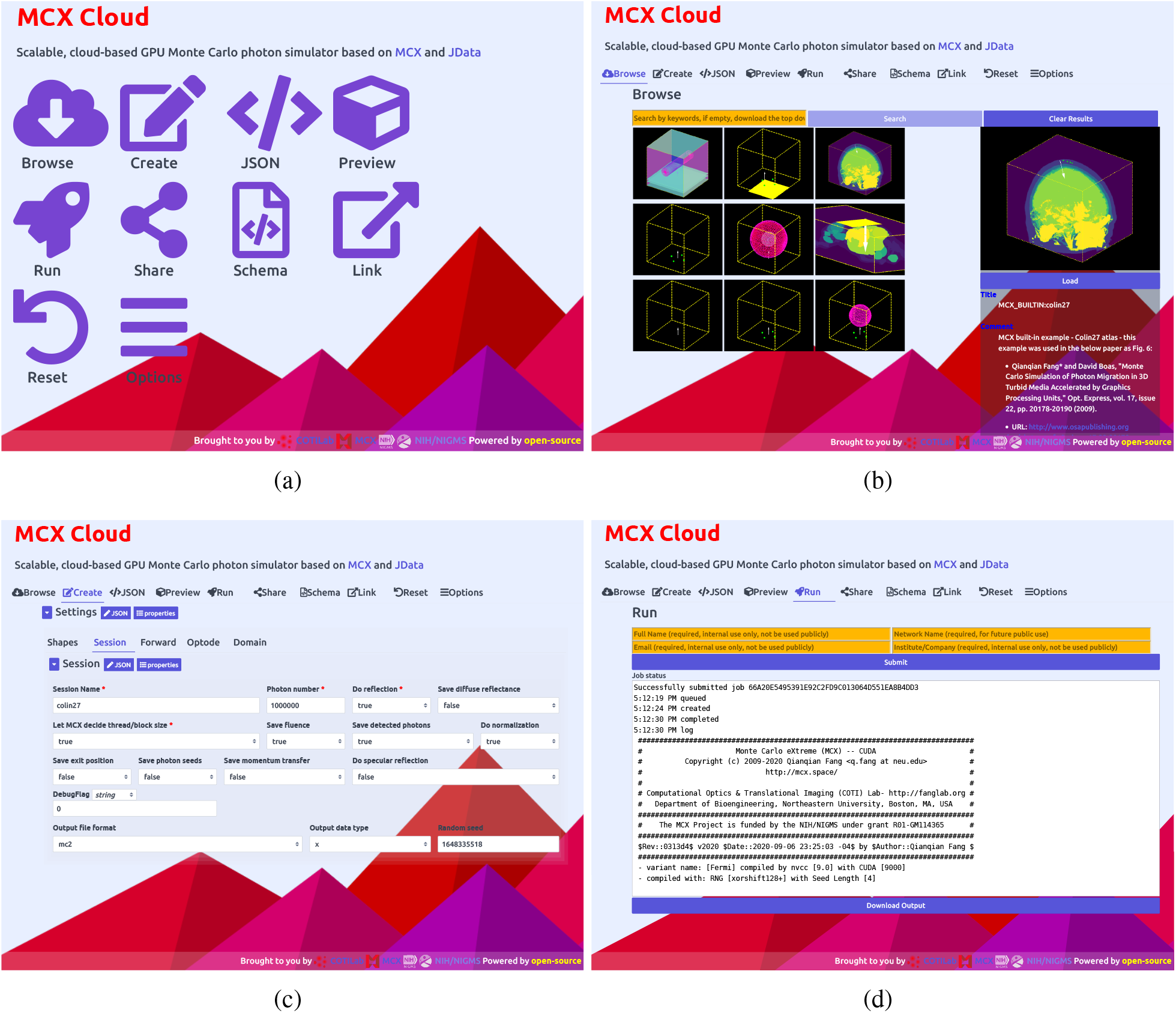
Sample screenshots of MCX Cloud graphical user interface (GUI) in a web browser, including (a) the main menu, (b) the “Browse” tab to download user-contributed simulations, (c) the “Create” tab for editing JSON-based input data validated by built-in schema, and (d) the “Run” tab to launch jobs to the cloud and monitor progress.

To show the 3-D domain rendering functions in the web GUI, in Fig. 3, we provide two screenshots showing both the shape-based and 3D-volume-based in-browser rendering via WebGL and Three.js APIs. The first rendering in Fig. 3(a) shows MCX’s built-in benchmark, “skinvessel”, which was derived from the benchmark used by mcxyz.^30^ The domain is described by JSON-based shape descriptors, consisting of 3 layers in the *z*-axis, a cylindrical object, and a disk-shaped source. In this screenshot, our front-end calls Three.js APIs to parse the shape descriptors and render each domain component in a “canvas” object. To give an example for rendering 3-D voxelated domain inputs, in Fig. 3(b), we show the web GUI rendering of the “digimouse” benchmark provided by MCX. The simulation domain is the segmented Digimouse atlas,^31^ described by a 190*×*496*×*104 unsigned-integer array with 21 tissue types. This 3-D segmented digital atlas is encoded in the JData N-D array format along with Zlib data compression (https://zlib.net) and Base64 encoding. The self-contained JSON input file is 188 kB in size. Using a WebGL rendering speed benchmark library, we have observed a speed of 180 to 300 frame-per-second (fps) for the “digimouse” on a range of desktop and laptop computers with dedicated NVIDIA GPUs; such speed drops to 20 to 60 fps when using this GUI on a laptop with Intel’s integrated GPUs.

**Fig 3.**
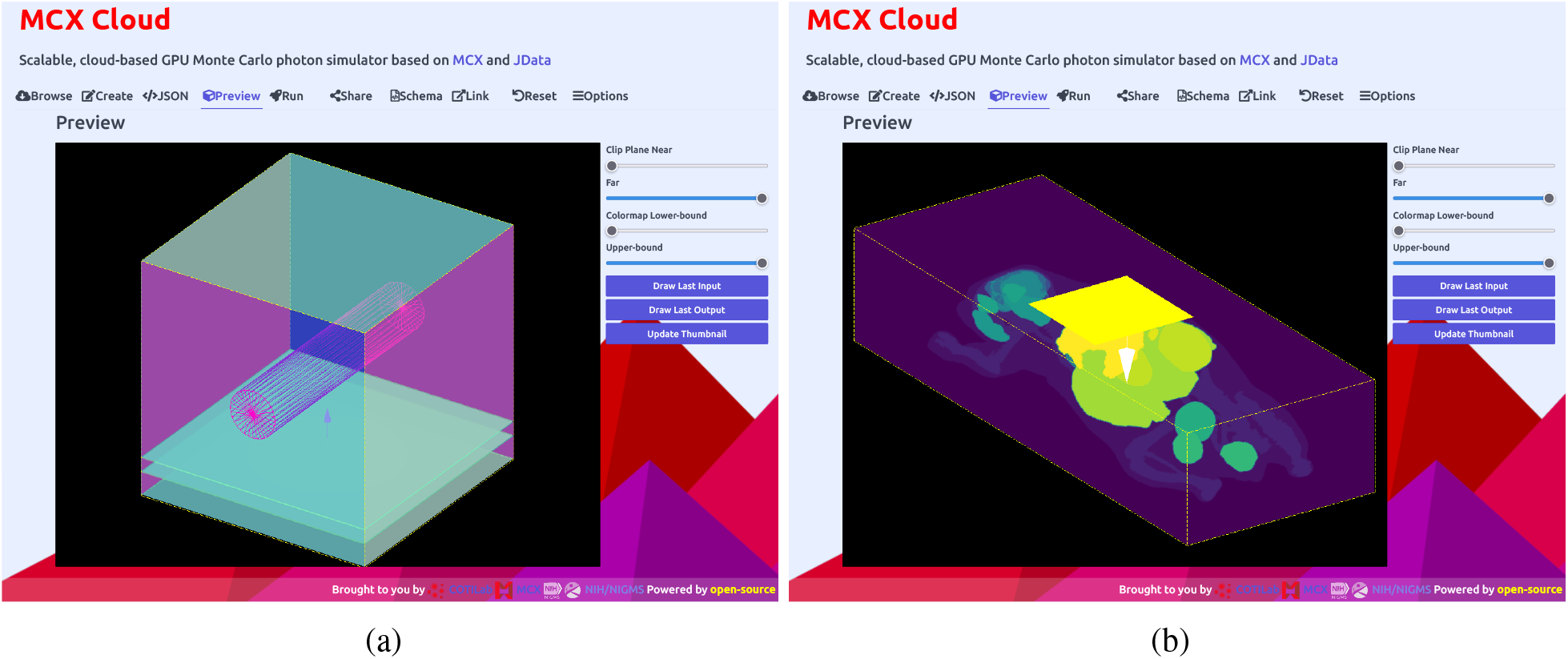
In-browser 3-D rendering samples of complex simulation domains, showing (a) the “skinvessel” benchmark and (b) the “digimouse” benchmark, using WebGL.

Our 3-D in-browser rendering tool also automatically renders MCX-computed fluence maps, also encoded in the JSON/JNIfTI format, returned by the server after the computation is completed. In Figs. 4(a) and 4(b), we show the 3-D views of the volumetric fluence rate (as MIP) obtained from the above two simulations. One can click on the “Download” button at the bottom of the rendering tab to download the entire 3-D output data file, encoded in the JSON/JNIfTI format, to the local disk for further analysis. Similarly, one can also click on the “Download” button in the “JSON” tab to download the web GUI generated JSON input file to his/her disk to locally run MCX on the user’s own computer.

**Fig 4.**
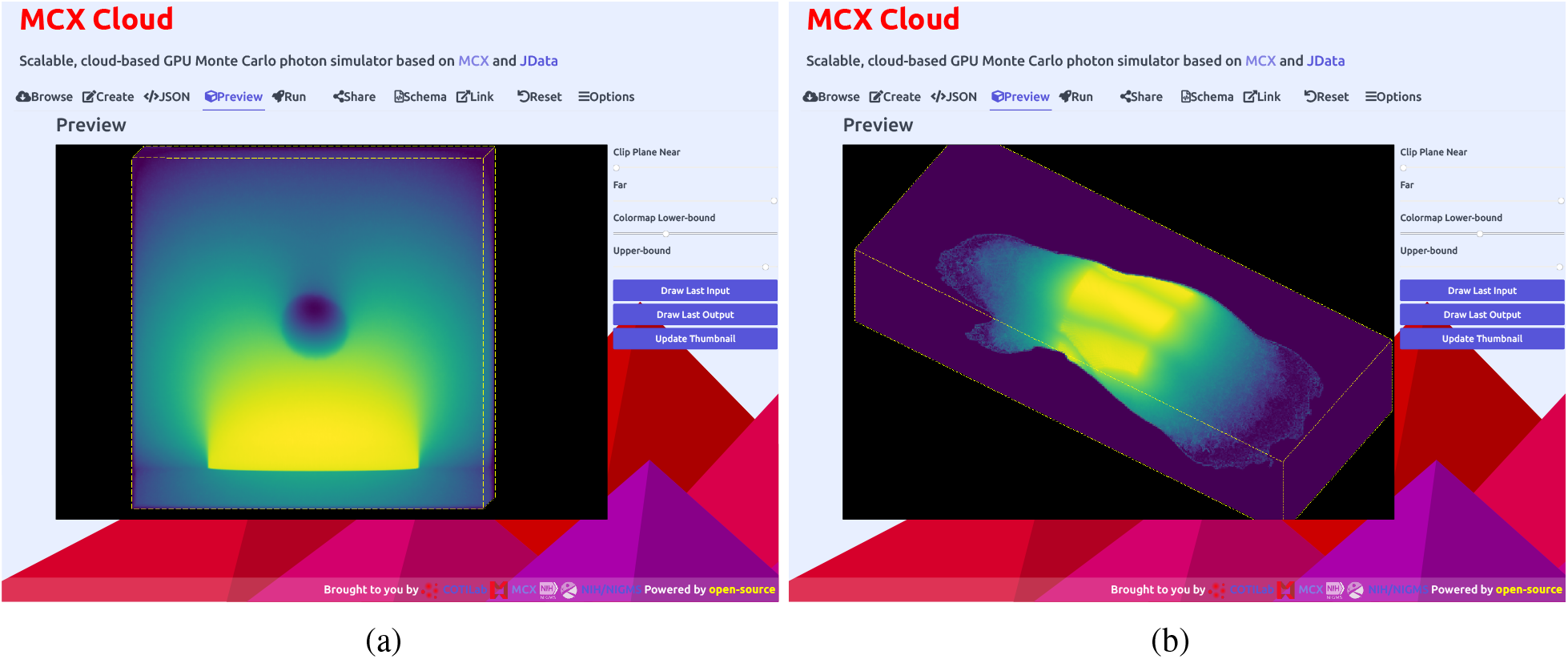
Volumetric rendering of the computed fluence-rate output from (a) the “skinvessel” benchmark and (b) the “digimouse” benchmark.

To demonstrate that one can use MCX Cloud to distribute a large simulation across multiple GPU devices installed in the Docker Swarm, we launch the “digimouse” benchmark simultaneously to 10 GPUs installed on the backend, each running 10^9^ photons, and record the elapsed time shown in a chart in Fig. 5. The overall simulation speed is 20,704 photon/ms if counting from the job submission time, or 21,834 photon/ms if counting from the start of the first job. This is about 3*×* of the average speed on all RTX 2080S nodes (6,775 photon/ms), and 9× of that on the GTX 1080 GPUs (2,374 photon/ms). We want to highlight that this sample simulation is designed to show the versatility of the platform without making any attempt to optimize to achieve maximum speed. The simulation speed can be easily improved by adjusting backend settings to increase the frequency of running the “mcxcloudd” server script and perform GPU-based load-balancing.

**Fig 5.**
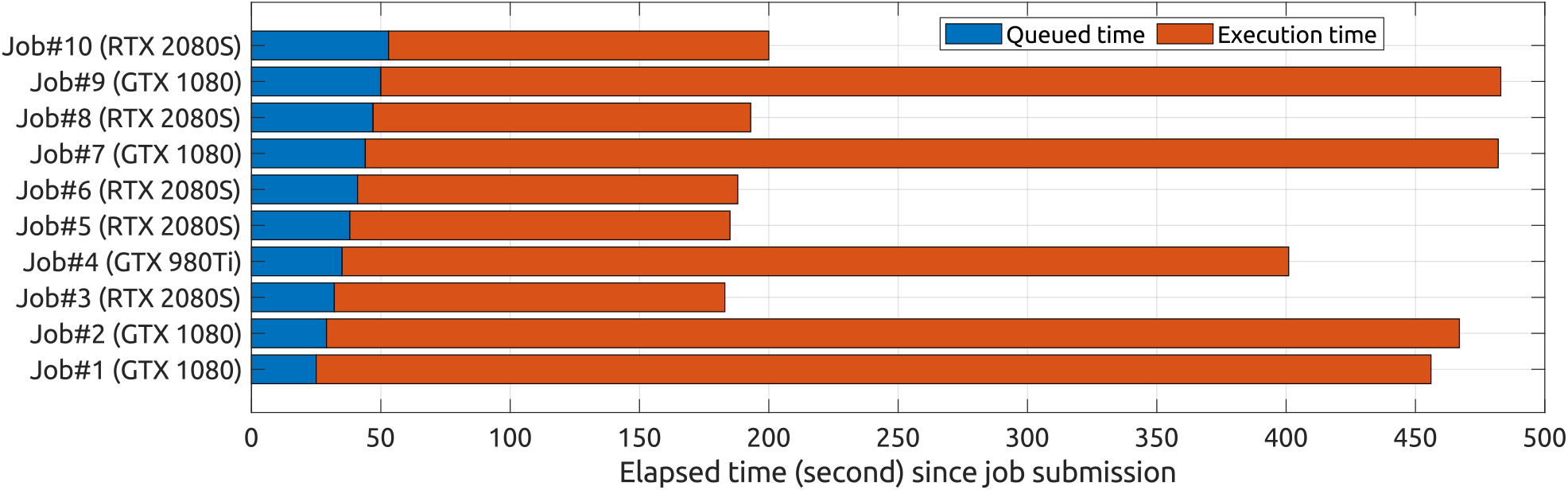
Elapsed time analysis for running the “digimouse” benchmark with a total of 10^10^ photons using 10*×* NVIDIA GPU devices via MCX Cloud. In this example, we used 5*×* RTX 2080SUPER, 4*×* GTX 1080 and 1*×* GTX 980Ti.

### 3.1 Discussions and Conclusion

Over the past decade, MC-based photon transport simulation has gained ample progress in terms of speed and accuracy in modeling increasingly complex anatomical structures. A list of free and open-source MC simulators with various levels of functionalities have been developed, published, and actively maintained by a number of research groups. While some of these open-source toolkits have successfully attracted a sizable user community, most of these tools were disseminated using a conventional download-and-install approach. In addition, many high-performance MC simulators require purchasing and installing high-end graphics cards on users’ own computers to maximize efficiency. For less-experienced users, properly configuring and using these specialized simulation tools can be key barriers.

This work specifically addresses challenges regarding the usability and availability of MC simulators as mentioned above. Particularly, we described an in-browser GPU-accelerated MC simulator and cloud-based service that can be launched anywhere a browser is available, including mobile devices such as a smartphone or a tablet. This system combines our decade-long, continual development in MCX light transport simulation software with state-of-the-art cloud-computing platforms, and offers a robust, scalable and forward-looking framework for a standardized, high-demand, high-throughput and community-focused MC modeling platform. Compared to the previously published online MC simulator, this new platform embraces the latest technologies in microservices, cloud-computing (containerization and orchestration), and web-based GUI design (AJAX, JSON, JSON Schema, jQuery, WebGL and Three.js), and demonstrates high flexibility and scalability that were not previously available.

We can not emphasize enough how adopting a standardized and web-friendly input/output data format in JSON/JData greatly simplified or even directly enabled the implementation of this lightweight yet highly versatile web-based platform. To be more specific, utilizing JSON to encode MCX’s input/output data allowed us to seamlessly integrate them with JavaScript and a web environment. Also, defining MCX’s input data using JSON schema allows the JSON Editor library to automatically create the JSON editing interface in our front-end. This in-browser JSON editor is not only intuitive to use, but also generates JSON data that automatically satisfies the specified schema. Similarly, adopting JSON and JData data annotations also allow MCX to store complex output data records, including volumetric fluence rate, partial-pathlengths, and various lightweight metadata in a unified, easy-to-read JSON format that can be readily transmitted, parsed and rendered inside a browser.

Although we use MCX at the backend to perform the underlying MC computation, our cloud computing system can be readily adapted to use any other MC simulators, as long as the alternative simulator also supports JSON/JData as the input/output data format and provides the corresponding JSON schema of the desired input JSON data structure (can be entirely different from those of MCX). For the same reason, our current web GUI can be directly used in combination with MCX-CL^5^ as the simulator in the backend if AMD or Intel GPUs are configured in Docker Swarm. This is because MCX-CL and MCX share nearly identical input/output formats. We are currently working on creating similar JSON/JData support for our MMC simulator,^10^ and anticipate that running MMC simulations on this cloud-computing platform will be supported in the near future.

From the benchmark results shown in Fig. 5, it is clear that this cloud computing platform can function not only as a parallel processor for simultaneously submitted jobs from multiple remote users, but also as a distributed high-performance computing platform to allow the running of a single simulation using all GPUs available. With more nodes and GPU devices added to the Docker Swarm, one should anticipate a nearly linear increase in the simulation speed when running large simulation loads.

Moving forwards, we aim to complete the migration of our MMC simulator^10^ to the JSON input/output data format, and make our web GUI readily usable for executing mesh-based MC simulations online. We will also focus on curating a comprehensive and reusable community-contributed MC simulation library and create standardized benchmarks to facilitate easy cross-validation between existing and emerging MC and diffusion solvers. In addition, we will monitor the utility of our GPU cloud and expand the capacity when necessary. We are also interested in upgrading the current Turing-/Pascal-based NVIDIA GPUs to the newer and more powerful generations as they become available to help users run their simulations in less time. We will release detailed tutorials and documentations on our MCX web site to guide users to configure and optimize their “private MCX cloud” when such guidance is necessary. In addition, containerization of MC simulators, like MCX, is only the beginning of building more sophisticated and automated bio-photonic data analysis pipelines. With more optical data analysis tools disseminated in a container environment, and more tools accepting the use of a standardized format, such as JSON/JData, as the input/output file format, the developers in our community will be able to create more sophisticated and automated data analysis processes using Docker compose, a standard tool to invoke multiple containerized applications.

The next step of our project also includes further solidification and dissemination of the Jdata specification (http://openjdata.org) for portable scientific data exchange, which has recently been funded by the NIH, including the exchange of volumetric data via the JNIfTI format,^28^ unstructured mesh data via the JMesh format^32^ etc. All of these JData-based data formats are fully JSON compatible and can be readily parsed by all existing JSON parsers and libraries. We strongly believe that providing such a universal data exchange platform permits all optical data analysis tools, and other scientific software in general, to efficiently share, exchange, integrate and automate hierarchical data records that are essential to scientific research. The convergence to a JSON-based data exchange platform also enables the research community to benefit from the latest NoSQL hierarchical database technology for large-volume and scalable scientific data storage and integration. Using MCX Cloud as a showcase, we sincerely invite all open-source MC simulator developers to consider supporting JSON-/JData-based data formats in their software to take advantage of these major benefits.

In summary, we report a highly scalable, easy-to-use, and cloud-computing-based in-browser MC simulation platform – MCX Cloud. This platform was built upon an array of modern open-source technologies, including the use of Docker containers and container orchestration to run GPU-based MC simulations across a robust, elastic, scalable, and distributed virtual GPU cluster. It also leverages the latest web-based technologies, such as JSON, JSON schema, AJAX, and We-bGL, to create an intuitive, easily expandable, and responsive web GUI. At the core of this cloud computing platform is our significantly improved MCX photon transport simulator, packaging numerous enhancements in GPU optimization and algorithmic features that we have developed and integrated over the past decade. We want to particularly highlight that this platform is fully open-source – we not only provide the source codes for the MCX simulator, but also those for the web GUI and server-side scripts – so that anyone can build a private cloud for internal use or modify these scripts to accommodate other similar solvers. In the meantime, we have built an initial GPU cloud containing 10*×* NVIDIA GPUs to help users execute MCX simulations without needing to purchase or maintain GPU hardware. Our online MCX simulation service is freely available at http://mcx.space/cloud.

## Disclosures

No conflicts of interest, financial or otherwise, are declared by the authors.

## Acknowledgments

This research is supported by the National Institutes of Health (NIH) grants R01-GM114365, R01- EB026998, and U24-NS124027. We would like to thank Leiming Yu and Yuhui Bao for their inputs and help in creating MCX Docker images.

**Qianqian Fang**, PhD, is currently an Associate Professor in the Bioengineering Department, Northeastern University, Boston, USA. He received his PhD degree from Thayer School of Engineering, Dartmouth College, in 2005. He then joined Massachusetts General Hospital and became an Instructor of Radiology in 2009 and Assistant Professor of Radiology in 2012, before he joined Northeastern University in 2015. His research interests include translational medical imaging devices, multi-modal imaging, image reconstruction algorithms, and high performance computing tools to facilitate the development of next-generation imaging platforms.

**Shijie Yan** is a doctoral candidate at Northeastern University. He received his BE degree from Southeast University, China, in 2013 and MS from Northeastern University in 2017. His research interests include Monte Carlo photon transport simulation algorithms, parallel computing, GPU programming and optimization.

## Notes

http://mcx.space/cloud

### Competing Interest Statement

The authors have declared no competing interest.

### Summary of Updates

Minor language updates following initial reviews

## References

1 L. V. Wang, S. L. Jacques, and L. Zheng, “MCML-Monte Carlo modeling of light transport in multi-layered tissues,” Comput. Methods Progr. Biomed. 47(2), 131–146 (1995).

2 E. Alerstam, T. Svensson, and S. Andersson-Engels, “Parallel computing with graphics processing units for high-speed Monte Carlo simulation of photon migration,” Journal of Biomedical Optics 13(6), 1 – 3 (2008).

3 Q. Fang and D. A. Boas, “Monte Carlo simulation of photon migration in 3D turbid media accelerated by graphics processing units,” Opt. Express 17(22), 20178–20190 (2009).

4 N. Ren, J. Liang, X. Qu, et al., “GPU-based Monte Carlo simulation for light propagation in complex heterogeneous tissues,” Opt. Express 18(7), 6811–6823 (2010).

5 L. Yu, F. Nina-Paravecino, D. Kaeli, et al., “Scalable and massively parallel Monte Carlo photon transport simulations for heterogeneous computing platforms,” J. Biomed. Opt. 23(1), 010504 (2018).

6 C. J. Zoller, A. Hohmann, F. Forschum, et al., “Parallelized Monte Carlo software to efficiently simulate the light propagation in arbitrarily shaped objects and aligned scattering media,” Journal of Biomedical Optics 23(6), 1–12 – 12 (2018).

7 T. Young-Schultz, S. Brown, L. Lilge, et al., “FullMonteCUDA: a fast, flexible, and accurate GPU-accelerated Monte Carlo simulator for light propagation in turbid media,” Biomed. Opt. Express 10, 4711–4726 (2019).

8 E. Margallo-Balbás and P. J. French, “Shape based Monte Carlo code for light transport in complex heterogeneous tissues,” Opt. Express 15(21), 14086–14098 (2007).

9 H. Shen and G. Wang, “A tetrahedron-based inhomogeneous Monte Carlo optical simulator,” Phys. Med. Biol. 55(4), 947–962 (2010).

10 Q. Fang, “Mesh-based Monte Carlo method using fast raytracing in Pluücker coordinates,” Biomed. Opt. Express 1(1), 165–175 (2010).

11 V. Periyasamy and M. Pramanik, “Monte Carlo simulation of light transport in turbid medium with embedded object—spherical, cylindrical, ellipsoidal, or cuboidal objects embedded within multilayered tissues,” Journal of Biomedical Optics 19(4), 1 – 10 (2014).

12 S. Yan, A. P. Tran, and Q. Fang, “Dual-grid mesh-based Monte Carlo algorithm for efficient photon transport simulations in complex three-dimensional media,” Journal of Biomedical Optics 24(2), 020503 (2019).

13 A. P. Tran and S. L. Jacques, “Modeling voxel-based Monte Carlo light transport with curved and oblique boundary surfaces,” Journal of Biomedical Optics 25(2), 1 – 13 (2020).

14 S. Yan and Q. Fang, “Hybrid mesh and voxel based Monte Carlo algorithm for accurate and efficient photon transport modeling in complex bio-tissues,” Biomed. Opt. Express 11, 6262–6270 (2020).

15 Y. Yuan, S. Yan, and Q. Fang, “Light transport modeling in highly complex tissues using the implicit mesh-based Monte Carlo algorithm,” Biomed. Opt. Express 12, 147–161 (2021).

16 D. Marti, R. N. N. Aasbjerg, P. E. E. Andersen, et al., “MCmatlab: an open-source, user-friendly, MATLAB-integrated three-dimensional Monte Carlo light transport solver with heat diffusion and tissue damage,” Journal of Biomedical Optics 23(12), 1 – 6 (2018).

17 A. A. Leino, A. Pulkkinen, and T. Tarvainen, “ValoMC: a Monte Carlo software and MATLAB toolbox for simulating light transport in biological tissue,” OSA Continuum 2, 957–972 (2019).

18 Q. Fang and S. Yan, “Graphics processing unit-accelerated mesh-based Monte Carlo photon transport simulations,” Journal of Biomedical Optics 24(11), 1 – 6 (2019).

19 J. Cassidy, A. Nouri, V. Betz, et al., “High-performance, robustly verified Monte Carlo simulation with FullMonte,” Journal of Biomedical Optics 23(8), 085001 (2018).

20 A. Doronin and I. Meglinski, “Online object oriented Monte Carlo computational tool for the needs of biomedical optics,” Biomed. Opt. Express 2(9), 2461–2469 (2011).

21 J. Joönsson and E. Berrocal, “Multi-Scattering software: part I: online accelerated Monte Carlo simulation of light transport through scattering media,” Opt. Express 28, 37612–37638 (2020).

22 W3C Working Group, “HTML 5 – A vocabulary and associated APIs for HTML and XHTML.” https://www.w3.org/TR/2008/WD-html5-20080122/ (2008).

23 Khronos WebGL Working Group, “WebGL 2.0 Specification.” https://www.khronos.org/registry/webgl/specs/latest/2.0/ (2017).

24 Qianqian Fang, “JData: A general-purpose data annotation and interchange format, Version 1.” https://github.com/fangq/jdata (2020).

25 A. Wright and H. Andrews and B. Hutton Eds, “JSON Schema: A Media Type for Describing JSON Documents.” https://json-schema.org/specification.html (2020).

26 R. Yao, X. Intes, and Q. Fang, “Direct approach to compute Jacobians for diffuse optical tomography using perturbation Monte Carlo-based photon ‘replay’,” Biomed. Opt. Express 9, 4588–4603 (2018).

27 R. Cox, “Official definition of the NIFTI1 header.” https://nifti.nimh.nih.gov/pub/dist/src/niftilib/nifti1.h (2007).

28 Qianqian Fang, “JNIfTI: An extensible file format for storage and interchange of neuroimaging data, Version 1.” https://github.com/fangq/jnifti (2020).

29 M. Wilkinson, M. Dumontier, I. J. Aalbersberg, et al., “The FAIR Guiding Principles for scientific data management and stewardship,” Scientific Data 3 (2016).

30 S. Jacques, “mcxyz software.” https://omlc.org/software/mc/mcxyz/.

31 B. Dogdas, D. Stout, A. F. Chatziioannou, et al., “Digimouse: a 3D whole body mouse atlas from CT and cryosection data,” Physics in Medicine and Biology 52, 577–587 (2007).

32 Qianqian Fang, “JMesh - A versatile data format for unstructured meshes and geometries, Version 1.” https://github.com/fangq/jmesh (2020).

